# Pan-sphingolipid profiling reveals differential functions of sphingolipid biosynthesis isozymes of *C. elegans*

**DOI:** 10.1101/2023.11.03.565432

**Authors:** Hui Luo, Xue Zhao, Zi-Dan Wang, Gang Wu, Yu Xia, Meng-Qiu Dong, Yan Ma

**Author notes:** Equal contribution (Hui Luo, Xue Zhao).

## Abstract

Multiple isozymes are encoded in the *C. elegans* genome for the various sphingolipid biosynthesis reactions, but the contributions of individual isozymes are characterized only in part. We developed a simple but effective reversed-phase liquid chromatography-tandem mass spectrometry (RPLC-MS/MS) method that enables simultaneous identification and quantification of ceramides (Cer), glucosylceramides (GlcCer), and sphingomyelins (SM), three important classes of sphingolipids from the same MS run. Validating this pan-sphingolipid profiling method, we show that nearly all 47 quantifiable sphingolipid species found in young adult worms were reduced upon RNA interference (RNAi) of *sptl-1* or *elo-5*, which are required for synthesis of the id17:1 sphingoid base. We also confirm that HYL-1 and HYL-2, but not LAGR-1, constitute the major ceramide synthase activity with different preference for fatty acid substrates, and that CGT-3 plays a greater role than CGT-1 does in producing glucosylceramides. Intriguingly, *lagr-1* RNAi lowers the abundance of all sphingomyelin species and that of several glucosylceramide species, which suggests that LAGR-1 may have functions beyond what is predicted. Additionally, RNAi of *sms-1, −2,* and *-3* all lower the abundance of sphingomyelins with an odd number of carbon atoms (mostly C21 and C23, with or without hydroxylation) in the N-acyl chain, and only *sms-1* RNAi does not elevate sphingomyelins containing even-numbered N-acyl chains. This suggests that sphingolipids containing even-numbered N-acyl chains could be regulated separately, sometimes in opposite directions, with those containing odd-numbered N-acyls, presumably monomethyl branched chain fatty acyls. We also find that ceramide levels are kept in balance with those of glucosylceramides and sphingomyelins.

## Introduction

Sphingolipids are among the most abundant classes of membrane lipids in eukaryotes, second only to glycerophospholipids (1,2). Distinct from glycerophospholipids, which are built from a glycerol-3-phosphate backbone, sphingolipids have a backbone of long-chain aliphatic amine, typically with zero or one double bond between C4 and C5, one amine group (C2), and a variable number of OH groups (C1, C3 and C4). (3). This sphingoid base is synthesized from long-chain fatty acyl-coenzyme A (CoA) and serine. In mammals, palmitoyl-CoA is often used as a substrate and the resulting sphingoid base is typically d18:1 (d denotes the two −OH groups, 18 is the total number of carbon atoms, and 1 is the number of double bond) (4).

Among three classes of sphingolipids—ceramides, glycosylceramides, and sphingomyelins—ceramides have the simplest structure with only a fatty acyl chain attached to the C2 amine group. Ceramides are synthesized on the endoplasmic reticulum (ER) membrane (5), and further transferring to the ceramide C1 hydroxyl group of sugar moieties on the *cis* and medial Golgi membrane (6,7) or of phosphocholine on the *trans* Golgi membrane (8) generates glycosylceramides or sphingomyelins, respectively.

As structural components of cell membranes, sphingomyelins are highly enriched in the outer leaflet of the plasma membrane (PM) (9), while ceramides and glycosylceramides are also present in high abundance in the endomembrane system in addition to the PM on both leaflets (8,10,11). By concentrating membrane receptors and signaling molecules, sphingolipid membrane microdomains—also known as lipid rafts—provide a central platform for efficient transduction of cellular signals (12). Sphingolipids and their metabolites also serve as second messengers to regulate cell growth, differentiation and cell death (11,13,14). Dysregulation of sphingolipid metabolism has been linked to many human diseases, including neurodegenerative conditions (15), Type II diabetes (16), cardiovascular diseases (17), and cancers (18).

Sphingolipids of the nematode *C. elegans* are rather peculiar compared to their mammalian counterparts (19,20). The sphingoid base of *C. elegans*, from which other worm sphingolipids such as ceramides, glycosylceramides, and sphingomyelins are derived, is predominantly id17:1 (83%), followed by a minor alternative id17:0 (12%) and trace amounts of others (21,20) (Fig. 1A). The two C17-iso-branched sphingoid bases are modified condensation products of serine and C15iso—a monomethyl branched chain fatty acid (mmBCFA) (22,23) (Fig. 1A). The glycosylceramides of *C. elegans* are found to be predominantly glucosylceramides and contain 2-hydroxylated N-acyl chains (19,24). For *C. elegans* ceramides and sphingomyelins, both hydroxylated and non-hydroxylated N-acyl chains are found (24). Like their mammalian counterparts, the N-acyl chains of worm sphingolipids are primarily saturated, long or very long chain fatty acids (25).

**Figure 1.**
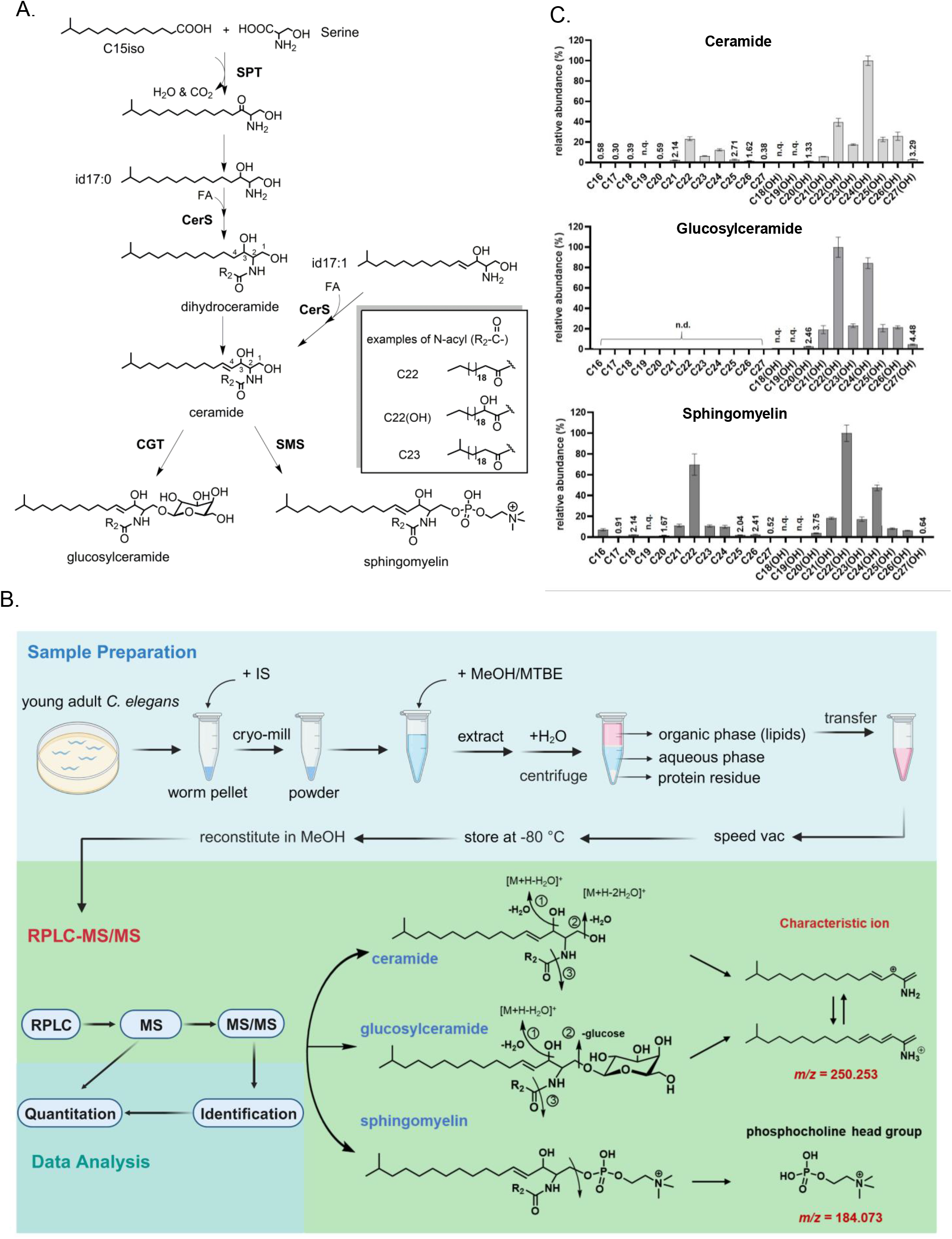
Establishment of a RPLC-MS/MS workflow for analysis of *C. elegans* sphingolipids. (A) A diagram of the *de novo* sphingolipid biosynthesis pathway in *C. elegans*. Serine palmitoyltransferase (SPT) catalyzes the condensation of a monomethyl branched chain fatty acid C15ISO and serine to form the precursor of the id17:1 sphingoid base. Ceramide synthase (CerS) catalyzes the acylation of a sphingoid base to form ceramide (Cer) or dihydroceramide. A ceramide glucosyltransferases (CGT) and a sphingomyelin synthases (SMS) transfers a glucose or a phosphocholine head group to the C1 hydroxyl of ceramide to form glucosylceramide (GlcCer) or sphingomyelin (SM), respectively. (B) The workflow for pan-sphingolipid profiling. Total lipids were extracted from 100 μL of tightly packed young adult *C. elegans.* The accurate mass of the precursor ion and that of the characteristic fragment ions were used to identify sphingolipids, *m/z* 250.253 for Cer and GlcCer, and *m/z* 184.073 for SM. (C) A total of 54 sphingolipids were identified from WT *C. elegans*. n.d. (not detected), n.q. (not quantifiable). The highest intensity species of each subclass was set to 100% relative abundance as a reference to scale other species of the same subclass.

Sphingolipids are essential for *C. elegans*; mutants that cannot synthesize sphingolipids arrest as L1 larvae and die (22,26–28). The underlying molecular mechanism has been unveiled through a series of investigations and glucosylceramide is found to be an essential growth signal that activates mTORC1 in *C. elegans* (27,29,30). Additionally, mutants lacking glucosylceramide fail to establish apicobasal polarity of the gut epithelial cells and suffer indigestion and larval arrest (10,29). The localization of membrane proteins can also be affected when glucosylceramide synthesis is disrupted (31). A specific C22-GlcCer is reported as a metabolite associated with longevity (32). Concerning other sphingolipids, ceramides are required for the anoxia response (33) as well as radiation-induced germline apoptosis in *C. elegans* (34), and sphingomyelins with very long-chain fatty acids (VLCFA) are implicated in the cuticle barrier function (35).

*C. elegans* worms cultured in the laboratory must synthesize sphingolipids on their own since their *E. coli* diet contains no sphingolipids (36,37). The *C. elegans* genome encodes a full set of enzymes required for biosynthesis of sphingolipids, including multiple isozymes whose functions are characterized incompletely. Specifically, there are three ceramide synthases (HYL-1, HYL-2, LAGR-1) (33,34,38,39), three UDP-glucose:ceramide glucosyltransferases (CGT-1, −2, and −3) (27) for the synthesis of glucosylceramides, and four sphingomyelin synthases (SMS-1, −2, −3, and −5). SMS-5 is a homolog of human SMS1, which is mainly localized to the Golgi where *de novo* synthesis of sphingomyelins takes place (40). SMS-1, −2, and −3 are homologs of human SMS2, which is localized to the Golgi membranes as well as the PM (8,40), where it can remodel sphingomyelins by resynthesizing them from ceramides generated by sphingomyelinases.

Except for HYL-1 and HYL-2 (33), the substrate specificity or product profiles of the above isozymes have not been characterized, as previous studies were concerned mainly with genetic analysis of developmental phenotypes (27,41).

In this study, we established a reversed-phase liquid chromatography-tandem mass spectrometry (RPLC-MS/MS) workflow for rapid and sensitive quantitation of *C. elegans* ceramides, glucosylceramides, and sphingomyelins all at once. In total, 54 sphingolipids were identified and 47 quantified from young adult worms. We validated this workflow using *C. elegans* worms in which the key enzymes responsible for generating the C15iso mmBCFA (*elo-5*) or the sphingoid base (*sptl-1*) were knocked down. Application of this workflow to isozyme characterization revealed rich information about the ceramide synthases, ceramide glucosyltransferases, and sphingomyelin synthases. For the ceramide synthases, *hyl-1* RNAi but not *hyl-2* or *lagr-1* RNAi reduced the abundance of all ceramide species. Regarding the ceramide glucosyltransferases, *cgt-3* RNAi but not *cgt-1* RNAi markedly decreased the abundance of all glucosylceramides quantified in *C. elegans*. Among the three worm homologs of human SMS2, none affected all SMs when knocked down, but each reduced the abundance of a subset of SMs. Our data also show that reduction of a sphingolipid subclass (or a subset of species within a subclass), as a direct result of RNAi of a targeted isozyme gene, is often accompanied by abundance changes of a different subclass (or other sphingolipids of the same class). This implicates a regulatory mechanism that can sense and respond to sphingolipid changes.

## Results

### A simple but effective LCMS-based workflow for analysis of *C. elegans* sphingolipids

We started out by optimizing the method for analyzing *C. elegans* sphingolipids. As shown in Fig. 1B, for lipid extraction from cryo-milled *C. elegans* samples, we chose methyl-*tert*-butyl ether (MTBE) (42) method for ease of handing. Alkali depletion of glycerolipids (24,43) is omitted because in our hands it introduced greater variations without improving sensitivity. For chromatographic separation of sphingolipids, we tried hydrophilic interaction liquid chromatography (HILIC), which separates lipids by their polar headgroups, and reversed-phase liquid chromatography (RPLC), which separates lipids by fatty acyl chains. We opted for RPLC for two reasons: first, the performance of RPLC is more consistent than HILIC; second, ceramides tend to have poorly shaped chromatographic peaks when separated on a HILIC column and are thus difficult to quantify. After optimization of the LC gradient, we finally arrived at a 15-min C18 RPLC run that enables separation of *C. elegans* sphingolipids (Methods and Sup Fig. S1).

From the high resolution, high mass accuracy MS/MS data, we only focused on sphingolipids built from the predominant id17:1 sphingoid base. For N-acyl chains, we considered fatty acids of 14–36 carbons with or without a hydroxyl group, and with or without a double bond after an initial consideration of up to four double bonds. Cyclopropane fatty acids, which the worms acquire from their bacterial food mainly in the form of C19Δ and C17Δ (23), were in effect also considered because they are of the same mass as C19:1 or C17:1. However, we did not find C19Δ, C17Δ, or any unsaturated N-acyls in *C. elegans* sphingolipids.

From wild-type *C. elegans* staged to adult day 1, we identified a total of 54 sphingolipid species including 22 Cers, 10 GlcCers, and 22 SMs (Sup Fig. S1), and 47 of them were quantifiable (Fig. 1C). As shown, *C. elegans* sphingolipids all have saturated N-acyl chains of 16–27 carbon atoms. The signal intensities of sphingolipids with hydroxylated N-acyl chains are much higher than those with non-hydroxylated N-acyl chains. In fact, all the glucosylceramides detected are hydroxylated. Our method cannot localize the hydroxyl group of the N-acyl, but based on the cumulative evidence provided by previous studies (19,44,45) it should be on C2 (Fig. 1A, inset, C22(OH) as an example). Among the ceramides, the highest MS signal intensity belonged to the C24(OH) species, followed by that of C22(OH). For glucosylceramides and sphingomyelins, the highest MS signal intensity species was found to be C22(OH) for both, followed by C24(OH) for glucosylceramides and C22 for sphingomyelins (Fig. 1C). The results above show that this simple, 15-min RPLC-MS/MS method enables effective profiling of *C. elegans* sphingolipids.

### Validation of the one-shot *C. elegans* sphingolipid analysis method

We validated our method on young adult *C. elegans* in which either the *elo-5* gene or the *sptl-1* gene was knocked down from the L1 stage by RNA interference (RNAi). SPTL-1 is the *C. elegans* ortholog of the catalytic subunit of human serine palmitoyltransferase (SPT), which is located on the ER (26,32). ELO-5 is a fatty acid elongase responsible for the biosynthesis of C15ISO (46,29), which is a substrate required for the first step of a chain of reactions that lead to the formation of the id17:1 sphingoid base (Fig. 1A, Fig. 2A, and Sup Fig. S2A). RNAi of either *sptl-1* or *elo-5* is expected to reduce the sphingolipid levels in *C. elegans*, and this is indeed the case (Fig. 2 and Sup Fig. S2). In *sptl-1* RNAi worms the total amount of Cer, GlcCer, and SM decreased by 94%, 49% and 23%, respectively (Fig. 2E-G). With respect to individual sphingolipid species, *sptl-1* RNAi resulted in a significant decrease for nearly all of them except for GlcCer C26(OH), SM C22, and SM C26(OH) (Fig. 2B-D). RNAi of *elo-5* markedly lowered the levels of all ceramide species and most of the SM species (Sup Fig. S2B and S2D). Intriguingly, glucosylceramides, which activate mTOR in *C. elegans* and are required for larval development (46,47), were least affected; only GlcCer C22(OH), the most abundant species in wild-type worms, had a significant, 66% decrease in *elo-5* RNAi worms (Sup Fig. S2C). This is in keeping with a recent study, which analyzed Cer and GlcCer but not SM, and showed that *elo-5* RNAi, decreased GlcCer C22(OH) but increased GlcCer C21(OH) and C23(OH) (48). This result invites a notion that under sphingoid base deficiency, *C. elegans* might channel what is available to synthesize GlcCer as much as possible to best relieve a life-threatening condition. Taken together, the *sptl-1* RNAi and the *elo-5* RNAi results validated this simple but effective pan-sphingolipid profiling method.

**Figure 2.**
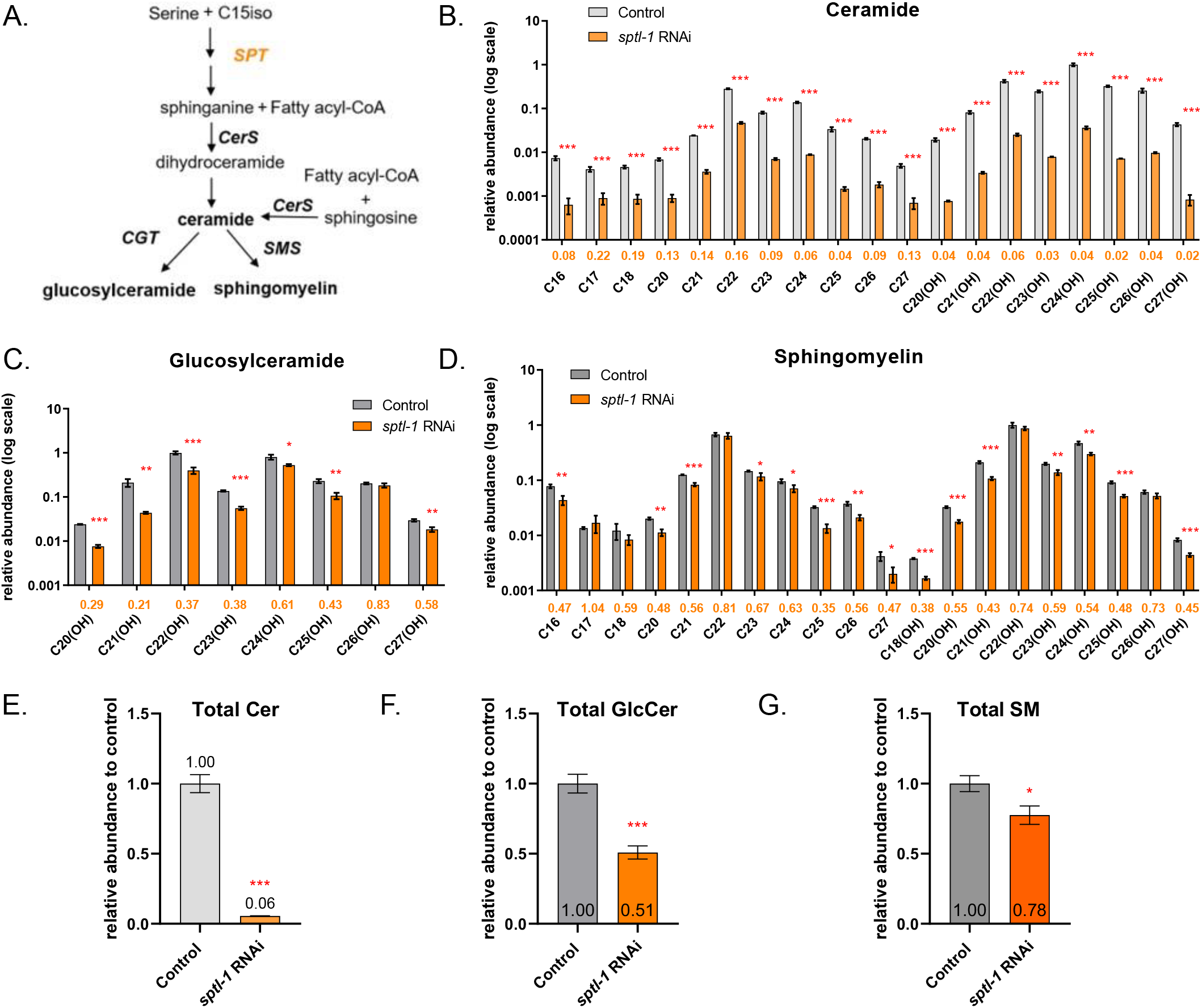
RNAi targeting the first key enzyme (SPT) in the *de novo* sphingolipid biosynthesis pathway caused a decrease of nearly all sphingolipids in *C. elegans*. (A) A simplified diagram of *de novo* sphingolipid biosynthesis in *C. elegans*. *sptl-1* encodes the catalytic subunit of SPT in *C. elegans*. (B-D) Relative abundance changes of individual Cer (B), GlcCer (C) and SM (D) species in *sptl-1* RNAi worms (log scale). *n* = 3 biological replicates, data are shown as mean ± standard deviation (∗, *P* < 0.05; ∗∗, *P* < 0.01; ∗∗∗, *P* < 0.001). The colored values below each bar graph indicate the fold change of abundance of the RNAi-treated group versus the control group. (E-G) Total Cer (E), GlcCer (F), and SM (G) level changes in *sptl-1* RNAi worms relative to the control RNAi worms.

### Distinct sphingolipid profiles of RNAi mutants lacking one of the three ceramide synthase isozymes

Ceramides are synthesized on the ER by ceramide synthases (CerS) out of a sphingoid base and fatty acyl CoA. (Fig. 1A and Fig. 2A) (11). *C. elegans* has three CerS isozymes, HYL-1, HYL-2, and LAGR-1 (49–51). Previous studies have found that loss-of-function*(lf)* mutants of the CerS genes exhibited different phenotypes. For example, *hyl-1(lf)* worms are resistant whereas *hyl-2(lf)* worms are sensitive to anoxia stress (33); *hyl-1(lf)* and *lagr-1(lf)* suppressed radiation-induced germ cell apoptosis whereas *hyl-2(lf)* did not (34); L1 larvae of *hyl-1(lf)* or *hyl-2(lf)*, but not of *lagr-1(lf)*, are more susceptible to death under starvation (52).

In this study we found that *hyl-1* RNAi lowered the abundance of all ceramide species, and this was most pronounced for those containing hydroxylated or non-hydroxylated C25∼C27 N-acyl chains (decreased by 49-76%, Fig. 3B-C, and 3F). Ceramides of C16∼C18 N-acyl chains also decreased to a similar extent upon *hyl-1* RNAi, but these species are of the lowest abundance in wild-type worms (Fig. 1C), so their decrease in absolute quantity is much less compared to the C25∼C27 species of high abundance.

**Figure 3.**
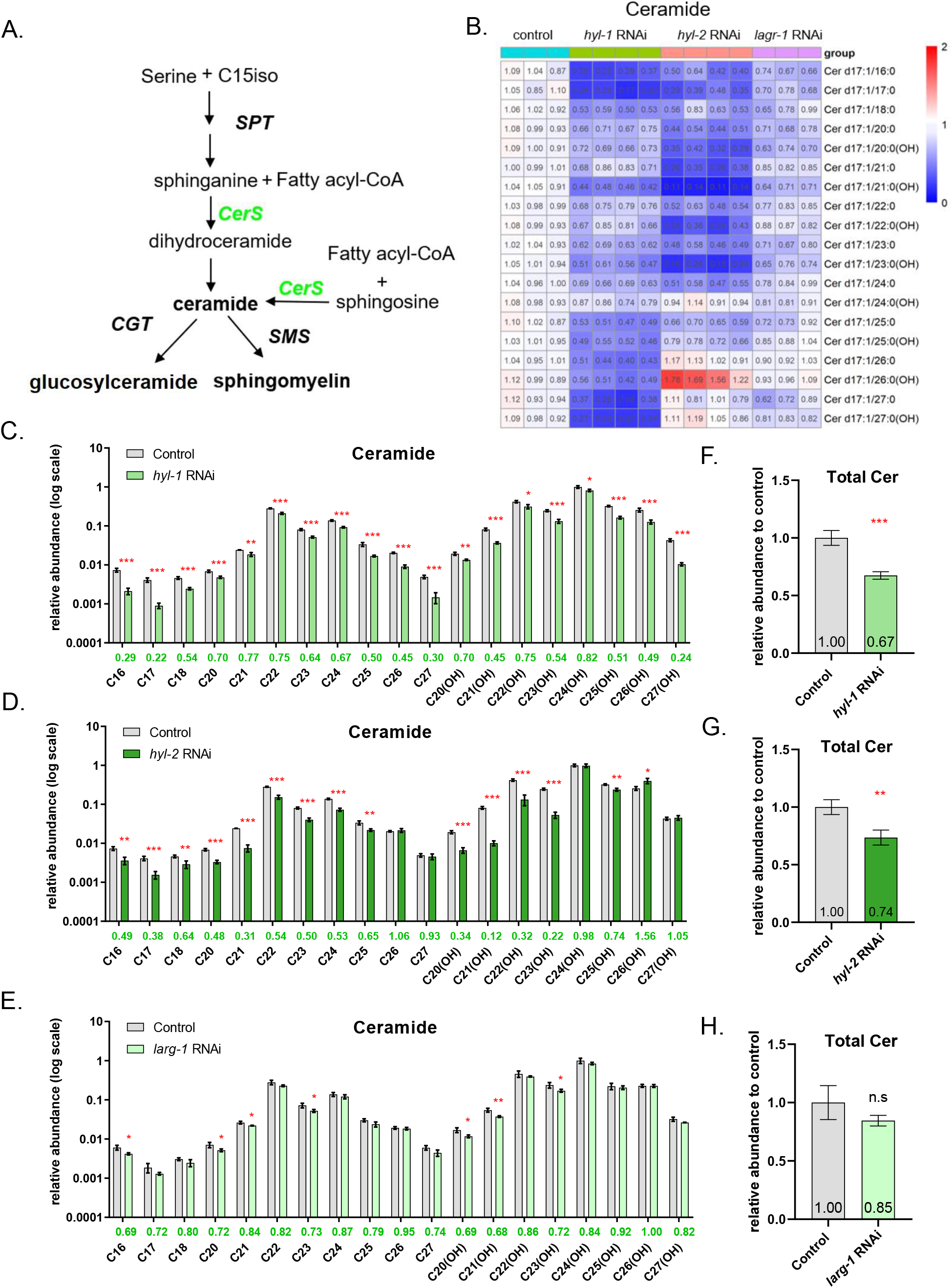
RNAi of each of the three ceramide synthase genes in *C. elegans* generated distinct ceramide profiles. (A) In the *de novo* sphingolipid biosynthesis pathway in *C. elegans*, CerS catalyzes the formation of ceramides. (B) A heatmap showing abundance changes of individual ceramide species in *hyl-1, hyl-2, lagr-1* RNAi worms relative to the control RNAi group. The average value of the three control RNAi samples, against which others were normalized, is set to 1. Abundance increase (fold change > 1) and decrease (fold change < 1) are indicated, respectively, by red and blue hues of varying saturation. (C-E) Relative abundance changes of individual ceramide species in *hyl-1* (C)*, hyl-2* (D)*, and lagr-1* (E) RNAi worms (log scale). *n* = 3 biological replicates, data are shown as mean ± standard deviation (∗, *P* < 0.05; ∗∗, *P* < 0.01; ∗∗∗, *P* < 0.001). The colored values below each bar graph indicate the fold change of abundance of the RNAi-treated group versus the control group. (F-H) Total Cer level changes in *hyl-1* (F)*, hyl-2* (G)*, and lagr-1* (H) RNAi worms relative to the control RNAi worms.

In contrast to *hyl-1* RNAi, *hyl-2* RNAi decreased most noticeably the amount of ceramide species containing hydroxylated C20∼C23 N-acyls by 46-88%. Ceramide species containing longer (C24∼27), hydroxylated N-acyl chains were not or only marginally decreased by *hyl-2* RNAi, among which Cer C26(OH) actually increased significantly by 56% (Fig. 3B, 3D, and 3G). These results are in agreement with the findings of an earlier study (33).

The sphingolipid profiles of the *lagr-1(lf)* mutant have not been analyzed before. Here we found that *lagr-1* RNAi had no or only marginal effect on ceramide levels (Fig. 3B, 3E, and 3H).

In addition to ceramides, pan-sphingolipid profiling allowed us to examine the effect of *hyl-1*, *hyl-2, and lagr-1* RNAi on GlcCer and SM (Sup Fig. S3). Aside from decreasing the levels of ceramide species of very long chain fatty acids (VLCFA, >24C), *hyl-1* RNAi also decreased the levels of GlcCer C27(OH) and five out of six VLCFA SM species, especially SM C27 and C27(OH) (Sup Fig. S3D and S3G).

In keeping with its effect on ceramides of hydroxylated or non-hydroxylated C20∼C23 N-acyls, *hyl-2* RNAi decreased the concentrations of four GlcCer species whose N-acyl chains varied from C20(OH) to C23(OH) (Sup Fig. S3A and S3C). Six out of eight SM species with hydroxylated or non-hydroxylated C20∼C23 N-acyl chains also decreased significantly in *hyl-2* RNAi worms (Sup Fig. S3E and S3G). Interestingly, *hyl-2* RNAi caused an increase of certain GlcCer and SM species, particularly the C24(OH) and C26(OH) varieties (Sup Fig. S3A, S3C, S3E and S3G).

Considering the modest effect of *lagr-1* RNAi on ceramides, with only 7 out of 19 Cer species displaying a weak but statistically significant decrease (by 16-32%), the effect of *lagr-1* RNAi on GlcCer and SM is disproportionally larger as it lowered the levels of 3 out of 8 GlcCer species (by 29-36%) and 13 out of 20 SM species (by 14%-37%) (Fig. 3 and Sup Fig. S3). Among the three ceramide synthase isozymes, *hyl-1* RNAi and *hyl-2* RNAi decreased the total amount of ceramides, *lagr-1* RNAi did not (Fig. 3F-H). Conversely, neither *hyl-1* RNAi nor *hyl-2* RNAi affected the total amount of GlcCer or SM (Sup Fig. S3B-D), whereas *lagr-1* RNAi unexpectedly decreased the total amount of SM and possibly that of GlcCer, too, although in the latter case the *p*-value (0.058) fell short of the 0.05 cutoff (Sup Fig. S3D and S3I).

### CGT-3 plays a greater role than CGT-1 does in producing glucosylceramides

*C. elegans* glucosylceramides all have hydroxylated N-acyls. Synthesis of glucosylceramides is catalyzed by UDP-glucose:ceramide glucosyltransferase (UGCG), for which *C. elegans* has three isozymes, CGT-1, −2, and −3. Mutations or RNAi treatments that compromise any single one of them did not show obvious phenotypes (27,53). However, *cgt-3;cgt-1* double mutant worms arrested and died at the L1 stage, *cgt-3;cgt-2* or *cgt-1;cgt-2* double mutants did not, and *cgt-2(lf)* did not further exacerbate the growth arrest phenotype of *cgt-3;cgt-1* worms (27,53). Although there is consensus on the growth arrest phenotype of *cgt* mutants, there is none when it comes to GlcCer. *Nomura et al.* reported a marked decrease of total GlcCer in the *cgt-1* or *cgt-3* single mutant (27), but *Marza et al.* found no such effect (53). Both studies used thin layer chromatography (TLC) to quantify the total GlcCer level. Here we analyzed individual GlcCer species by high-resolution mass spectrometry and focused on *cgt-1* and *cgt-3* (Fig. 4 and Sup Fig. S4).

**Figure 4.**
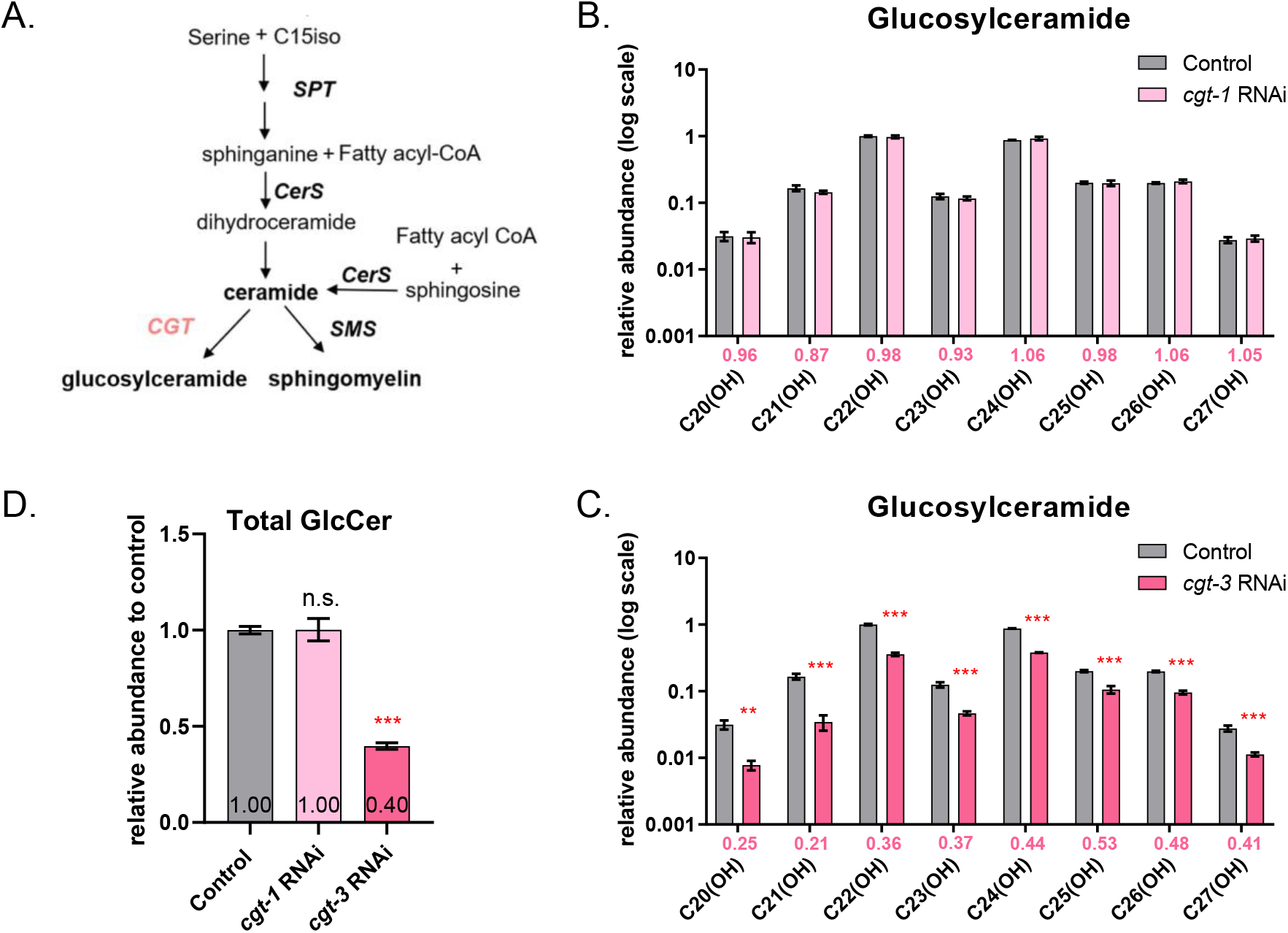
RNAi of *cgt-3* but not *cgt-1* decreased GlcCer levels. (A) In the *C. elegans* sphingolipid biosynthesis pathway, CGT catalyzes the formation of glucosylceramides. (B-C) Relative abundance changes of individual GlcCer species in *cgt-1* (B) *and cgt-3* (C) RNAi worms. *n* = 3 biological replicates, data are shown as mean ± standard deviation (∗, *P* < 0.05; ∗∗, *P* < 0.01; ∗∗∗, *P* < 0.001). The colored values below each bar graph indicate the fold change of abundance of the RNAi-treated group versus the control group. (D) Total GlcCer levels in *cgt-1* or *cgt-3* RNAi worms relative to that in control RNAi worms.

We found that *cgt-1* RNAi had no effect on GlcCer levels (Fig. 4B), and neither did it have a sizable effect on ceramide or sphingomyelin levels (Sup Fig. S4A, C, D, and F). In contrast, *cgt-3* RNAi reduced the total GlcCer level and every single GlcCer species quantified by 47-64% (Fig. 4C-D). Additionally, *cgt-3* RNAi caused a decrease of all the ceramide species with hydroxylated N-acyl chains by 37-57% (Sup Fig. S4B-C). In other words, when the activity of CGT-3 is compromised, both the substrate (Cer with 2-OH N-acyl chain) and the product (GlcCer with 2-OH N-acyl chain) are reduced, instead of a product decrease leading to a substrate buildup. This could suggest a feedback mechanism that keeps the substrate and the product levels in balance. Sphingomyelins were not affected by *cgt-3* RNAi in a marked way (Sup Fig. S4E-F).

### Overlapping sphingolipid profiles of RNAi mutants lacking a sphingomyelin synthase SMS −1, −2, or −3

*C. elegans* sphingomyelin synthases SMS-1, −2 and −3 are homologs of human SMS2, which is located at the PM as well as the Golgi. Human SMS2 is a membrane protein and its catalytic site faces the extracellular space or the Golgi lumen (40,54). Sphingomyelins are synthesized *de novo* in the Golgi, and those in the outer leaflet of the PM could be remodeled through the actions of acid sphingomyelinases and sphingomyelin synthases. Much is unknown about the functions of *C. elegans* SMSs other than that *sms-1(lf)* suppressed clozapine-induced developmental delay or lethality, whereas *sms-2(lf)* or *sms-3(lf)* did not (55).

In this study, we found that *sms-2* and *sms-3* RNAi worms have similar sphingomyelin profiles, which are different from that of *sms-1* RNAi worms (Fig. 5). As shown, *sms-1* RNAi caused a 30-52% decrease of sphingomyelin species whose N-acyl groups, hydroxylated or not, contain an odd number of carbon atoms such as C21, C23, or C25 (Fig. 5B and Sup Fig. S5G), which are presumably mmBCFAs. One even-numbered N-acyl sphingomyelin species (SM C24) decreased by 39% in *sms-1* RNAi worms but it was not statistically significant (Fig. 5B-C). For *sms-2* RNAi and *sms-3* RNAi, although they lowered the quantities of four sphingomyelin species containing an N-acyl group of odd numbered carbons (C21, C23 and their hydroxylated forms)—somewhat similar to *sms-1* RNAi—they elevated the levels of several SM species with even-numbered N-acyl chains such as C22, C24(OH), and C26(OH), which was not evident under *sms-1* RNAi. *sms-3* RNAi additionally increased SM C16, SM C18, and SM C20(OH) (Sup Fig. S5H, I). The observed increase of the sphingomyelin species containing certain even-numbered N-acyl chains could not be predicted. In brief, our data suggest that *sms-1, −2,* and *-3* all contribute to the production of sphingomyelin species containing a C21, C23, C21(OH), or C23(OH) N-acyl group (Fig. 5), which are of medium intensity (Fig. 1C). Further studies are needed to find out whether SMS-1, −2, and −3 are redundant for producing high-abundance sphingomyelins with even-numbered N-acyls, or whether a different sphingomyelin synthase is responsible for this function.

**Figure 5.**
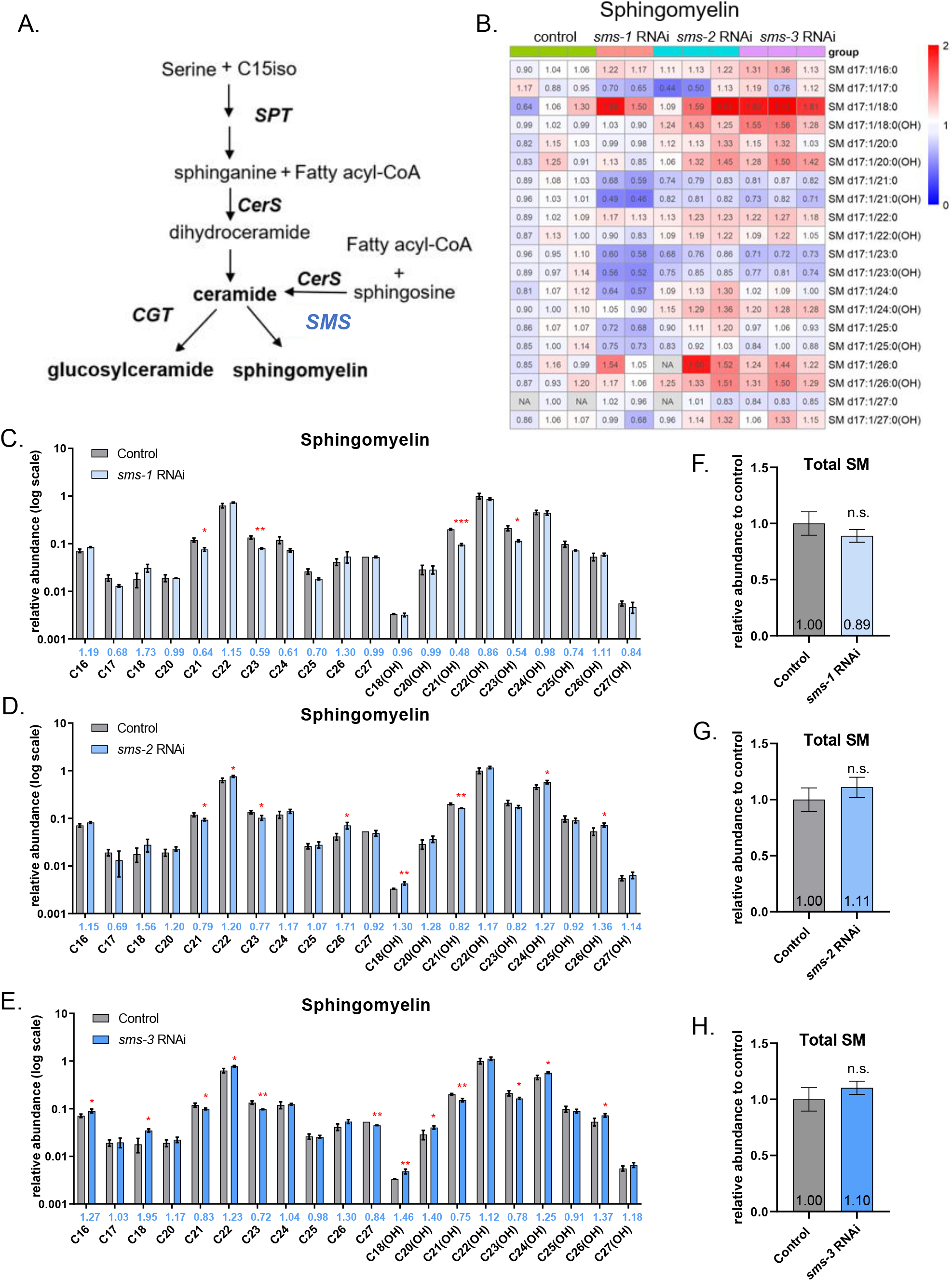
RNAi of *sms-2* or *sms-3* generated similar SM profiles, which differed from that by *sms-1* RNAi. (A) In the *C. elegans* sphingolipid biosynthesis pathway, SMS catalyzes the formation of sphingomyelins. (B) A heatmap showing abundance changes of individual sphingomyelin species in *sms-1, sms-2, sms-3* RNAi worms relative to the control RNAi group. The average value of the three control RNAi samples, against which others were normalized, is set to 1. Abundance increase (fold change > 1) and decrease (fold change < 1) are indicated, respectively, by red and blue hues of varying saturation. (C-E) Relative abundance changes of individual SM species in *sms-1* (C)*, sms-2* (D)*, and sms-3* (E) RNAi worms (log scale). *n* = 3 biological replicates, data are shown as mean ± standard deviation (∗, *P* < 0.05; ∗∗, *P* < 0.01; ∗∗∗, *P* < 0.001). The colored values below each bar graph indicate the fold change of abundance of the RNAi-treated group versus the control group. (F-H) Total SM levels in *sms-1* (F)*, sms-2* (G)*, and sms-3* (H) RNAi worms relative to the control RNAi worms.

Displayed in Sup Fig. S5 are four additional findings from the pan-sphingolipid profiling experiments. (1) *sms-3* RNAi reduced the total amount of ceramides (Sup Fig. S5E). (2) *sms-2* RNAi and *sms-3* RNAi, but not *sms-1* RNAi lowered the levels of nearly all ceramide species whose N-acyl groups containing an odd number of carbon atoms, hydroxylated or not (Fig. 5A and Sup Fig. S5A-C). (3) *sms-2* RNAi and *sms-3* RNAi both lowered the abundance of GlcCer C21(OH) (Sup Fig. S5E and 5H-I). (4) *sms-1* RNAi increased the levels of three GlcCer species C24(OH), C26(OH), and C27(OH) (Sup Fig. S5E and S5G and Fig. 5C).

## Discussion

For a biosynthetic pathway of the simplest type, one would expect that following the inactivation of an enzyme, the product of the reaction catalyzed by that enzyme as well as other metabolic products further downstream will deplete, while the substrate and other precursor metabolites further upstream will accumulate. This is not the case for sphingolipid biosynthesis as can be seen from the sphingolipid profiles of *cgt-3* RNAi worms and *sms-2* or *sms-3* RNAi worms. The decrease of GlcCer as a direct consequence of *cgt-3* RNAi was accompanied not by an increase but a decrease of the ceramide substrate (Fig. 4 and Sup Fig. S4). The lack of an increase could not be explained by surplus ceramides being used to make more sphingomyelins, as the total sphingomyelin level hardly changed despite small increase or decrease of certain species (Sup Fig. S4). Likewise, under *sms-2* or *sms-3* RNAi, the decrease of SM C21/23 and SM C21/23(OH) was accompanied not by an increase but a decrease of the ceramide substrate (Fig. 5 and Sup Fig. S5). Again, the lack of an increase was not because surplus ceramides were used to make more GlcCer (Sup Fig. S5). Hence, the sphingolipid biosynthesis pathway is not a simple one; it appears to be under strict regulation to prevent an overabundance of ceramides when a downstream biosynthesis step is obstructed. In other words, the ceramide levels need to be kept in balance with those of GlcCer and sphingomyelins.

The sphingolipid profiles of the *sms-2* and *sms-3* RNAi worms revealed an interesting phenomenon (Fig. 5). The decrease of SMs C21, C21(OH), C23, and C23(OH) was accompanied by an increase of SMs C22, C24(OH), and C26(OH). Similarly, in *elo-5* RNAi worms, the decrease of GlcCer C22(OH) was accompanied by an increase of GlcCer C21(OH) and GlcCer C23(OH) (Sup Fig. S2). This suggests that there may be a mechanism that could up-regulate certain sphingolipids of even-numbered N-acyl chains to compensate a decrease of sphingolipids of odd-numbered N-acyls, and vice versa. We have noticed that across the RNAi samples examined in this study, the abundance changes of GlcCer C23(OH), SM C21, and SM C21(OH) anti-correlated with most other sphingolipids including 16 out 21 species with even-numbered N-acyls (Sup Fig. S6). In keeping with this, the abundance changes of GlcCer C26(OH) and SM C26(OH) anti-correlated with Cer C21/C21(OH)/C23(OH), GlcCer C23(OH), and SM C21/C23/C21(OH), all with odd-numbered N-acyls (Sup Fig. S6). We thus propose that sphingolipids of even-numbered N-acyls may be regulated separately—at times oppositely—from those of odd-numbered, presumably mmBCFA N-acyls.

In this study, we identified from young adult *C. elegans* a total of 54 species out of three classes of sphingolipids Cer, GlcCer, and SM. In Supplemental Table S1, the sphingolipid identification results from this study are compared with those from previous ones. It is evident that the high-abundance Cer, GlcCer, and SM species were identified consistently across studies. The absence of GlcCer species with non-hydroxylated N-acyl chains, first reported in 1995 (19), was later verified by four studies including this one (39,43,24) and countered by one study (25). This comparison shows that the pan-sphingolipid profiling method developed here is sensitive and accurate, and it enables comprehensive analysis of *C. elegans* Cer, GlcCer, and SM.

## Materials and methods

### Worm culture and sample collection

The wild type N2 worms were fed with *E. coli* OP50 on nematode growth medium (NGM) plates and cultured at 20°C (56). To obtain synchronized animals, gravid adults were washed off plates with M9 buffer, then bleached with 30% sodium hypochlorite containing 0.75 M KOH to obtain the eggs. After hatching, the synchronized L1 worms were seeded onto fresh RNAi plates until harvest at young adult stage. Worms were washed three times with M9 buffer and pellet on ice, a total volume of 100 μl worm pellet was collected for each sample. For each RNAi conditions, three biological replicates were collected unless otherwise indicated.

### RNAi

RNAi assays were performed at 20°C using the feeding method as previously described (57). The *Escherichia coli* strain HT115 transfected with L4440 (empty vector) was used as control. The other RNAi bacterial strains were obtained from the Ahringer RNAi library or the Vidal library. All RNAi clones were verified by sequencing. Efficacy of RNAi was verified by growth or lifespan phenotypes when available. For example, *sptl-1* or *elo-5* RNAi worms die within days after reaching adulthood.

### Lipid extraction from *C. elegans*

The methyl-tert-butyl ether (MTBE)/methanol method was used for lipid extraction (42). After adding a fixed amount (0.3 μmol for each) of the internal standards (SM d18:1/12:0, Cer d18:1.10:0, GlcCer d18:1/12:0), worms were cryo-milled (Retch MM400). 225 μl methanol and 750 μl MTBE were added to the sample, incubate at 4 °C for 10 min. 200 μl water was added and the mixture was centrifuged at 4000 g for 8 min. The upper MTBE layer were collected and dried in the speed vacuum concentration system (Speed Vac). The extract was stored at −80°C and resolved in MeOH before analysis.

### LC-MS/MS analysis

LC-MS/MS analysis of sphingolipids was performed on a Thermo Vanquish UHPLC coupled with a Thermo Q Exactive HF-X hybrid quadrupole-Orbitrap mass spectrometer in positive ESI mode. The LC method was adapted from a published assay (58). Separation was carried out by a Waters ACQUITY UPLC BEH C18 column (1.7 μm, 2.1[×[100[mm). Column temperature was 60 °C. Mobile phases consisted of water (A) and ACN: IPA 4:3 (B), both with 10 mM ammonium acetate and 0.1% formic acid. The following gradient was applied at a flow rate of 0.5 mL/min: 85% B (0-1 min), 85-100% B (1-3 min), 100% B (3-8 min), 100-85% (8-12 min), 85% B (12-15 min). Samples were resuspended with 200 µL methanol and 2 µl was injected.

Full-scan mass spectra were acquired in the range of *m/z* 100 to 1200 with the following ESI source settings: spray voltage: 3.5 kV, auxiliary gas heater temperature: 380 °C, capillary temperature: 320 °C, sheath gas flow rate: 30 units, auxiliary gas flow: 10 units. MS1 scan parameters included resolution 60000, AGC target 3e6, and maximum injection time 200 ms. MS2 scan parameters included resolution 30000, AGC target 2e5, and maximum injection time 100 ms. Normalized collision energy was 30.

### Data processing and statistical analysis

Data preprocessing, including peak picking and alignment, was performed using MS-DIAL (59) software (v 4.16). An in-silico MS/MS library was developed for the annotation of ceramide (Cer), sphingomyelin (SM) and glucosylceramide (GlcCer). First, theoretical *m/z* of sphingolipids with d17:1 sphingoid base and fatty acyls from C14 to C36 were calculated. Hydroxylated or unsaturated species were also included. After that, the fragmentation patterns of Cer, SM and GluCer were investigated from the experimental MS/MS spectra of reference standard compounds, e.g., Cer d18:1/18:0, HexCer d18:1/15:0 and SM d18:1/18:0. Here the d18:1 species were used instead of d17:1 ones as the latter were not commercially available. Characteristic product ions and their relative abundances were used to predict the MS/MS spectra of d17:1 sphingolipids. For example, fragmentation of Cer d18:1/18:0 [M+H-H_2_O]^+^ yielded product ions at *m/z* 282.279 (C_18_H_36_NO^+^, 10% intensity of the base peak), 264.269 (C_18_H_34_N^+^, 100%), and 252.269 (C_17_H_34_N^+^, 10%), which originated from the C18 sphingosine (C_18_H_37_NO_2_). Therefore, replacing the C18 sphingosine with the C17 one gave the product ions of d17:1 Cer at *m/z* 268.264 (C_17_H_34_NO^+^, 10%), 250.253 (C_17_H_32_N^+^, 100%) and 238.253 (C_16_H_32_N^+^, 10%). Fragmentation pattern of HexCer was basically the same as Cer. For SM, the only product ion was phosphorylcholine headgroup at *m/z* 184.073, regardless of their sphingosine and fatty acyl compositions. These fragment ions were combined with the calculated precursor *m/z* to generate the in-silico library in MSP format using an in-house Java program. The MSP file was then converted to NIST lib format and searched against the experimental data with NIST MSPepSearch software. Hits with dot product score >600 were kept.

For quantitation, peak heights of the annotated sphingolipids were normalized with their corresponding internal standards. To compare the differences in each sphingolipid species between the RNAi group and the control group, Student’s t test was performed. *, *p* < 0.05; **, *p* < 0.01; ***, *p* < 0.001; ns, *p* ≥ 0.5.

## Data availability

The raw lipidomics data from HPLC/MS-MS measurements are available upon request.

## Acknowledgement

The authors gratefully acknowledge financial support from the Ministry of Science and Technology of China (2020YFF01014505 to Y.M and M.Q.D.), National Natural Science Foundation of China (NSFC-ISF 32061143020 to M.Q.D.), National Natural Science Foundation of China (NSFC 22225404 to Y.X.) the municipal government of Beijing (in the form of NIBS intramural grants), TIMBR, and Tsinghua University.

## CRediT author statement

H.L., M.Q.D., Y.X., X.Z. conceptualization; Y.M., Y.X., X.Z. methodology; H.L., Y.M., X.Z., Z.D.W., G.W., investigation, H.L., M.Q.D., Y.M. writing – original draft; H.L., M.Q.D., Y.X., Y.M., Z.D.W., X.Z. writing –review & editing; H.L., X.Z., Y.M. visualization; Y.X., M.Q.D., Y.M. supervision; Y.X., M.Q.D., Y.M. project administration; Y.X., M.Q.D., Y.M. funding acquisition.

