## supplemental figures for "Pan-sphingolipid profiling reveals differential functions of sphingolipid biosynthesis isozymes of *C. elegans*"

**Supplemental Figure S1. Separation of *C. elegans* sphingolipids on a C18 reverse phase column.**

(A) The retention time as a function of the number of carbon atoms in the N-acyl chain for each of the 54 sphingolipids identified from wild-type *C. elegans*. All contain an id17:1 sphingoid base and a saturated N-acyl chain varying from C16 to C27. (B) Representative chromatographic peaks of selected Cer, GlcCer, or SM species found in *C. elegans*. Colored Blue are three d18:1 sphingolipids used as internal standards, which are set to 100% relative abundance to scale the intensities of worm sphingolipids.

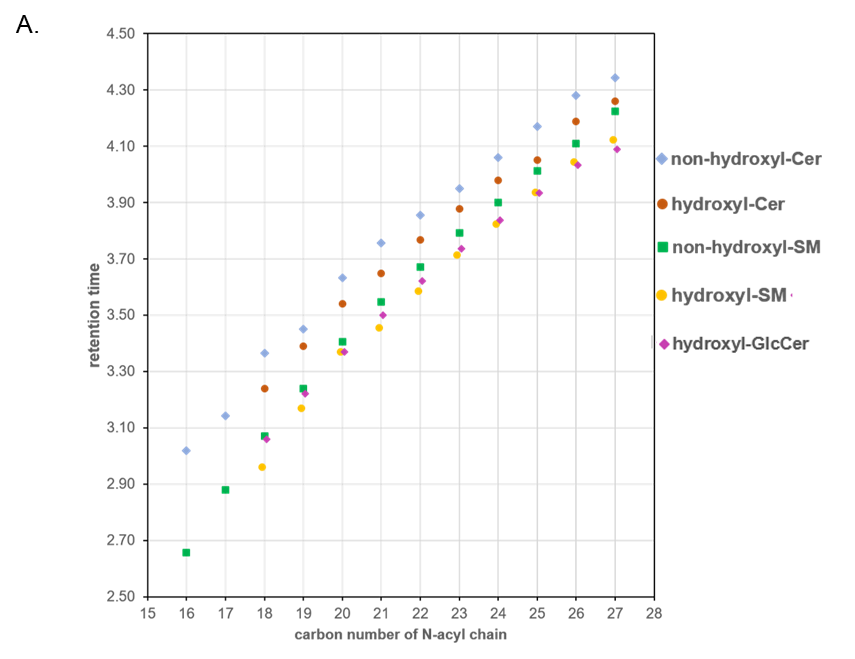

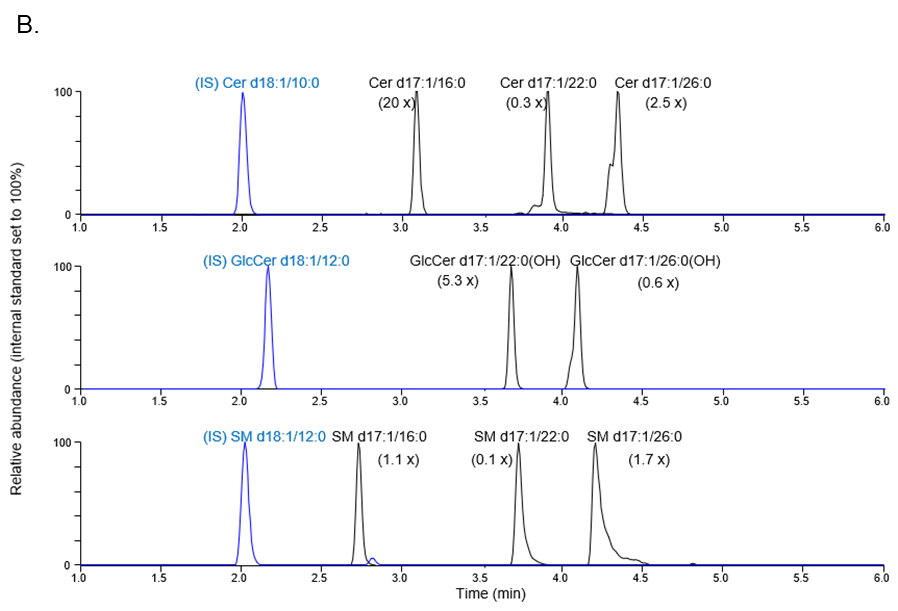

**Supplemental Figure S2. RNAi of *elo-5*, a long-chain mmBCFA elongase gene, caused an abundance decrease for all but a few sphingolipids in *C. elegans***

(A) ELO-5 catalyzes the formation of C15ISO, a substrate required for the synthesis of the sphingoid base. (B-D) Relative abundance changes of individual Cer (B), GlcCer (C) and SM (D) species in *elo-5* RNAi worms (log scale). *n* = 3 biological replicates, data are shown as mean ± standard deviation (∗, *P* < 0.05; ∗∗, *P* < 0.01; ∗∗∗, *P* < 0.001). The colored values below each bar graph indicate the fold change of abundance of the RNAi-treated group versus the control group. (E-G) Total Cer (E), GlcCer (F), and SM (G) level changes in *elo-5* RNAi worms relative to the control RNAi worms.

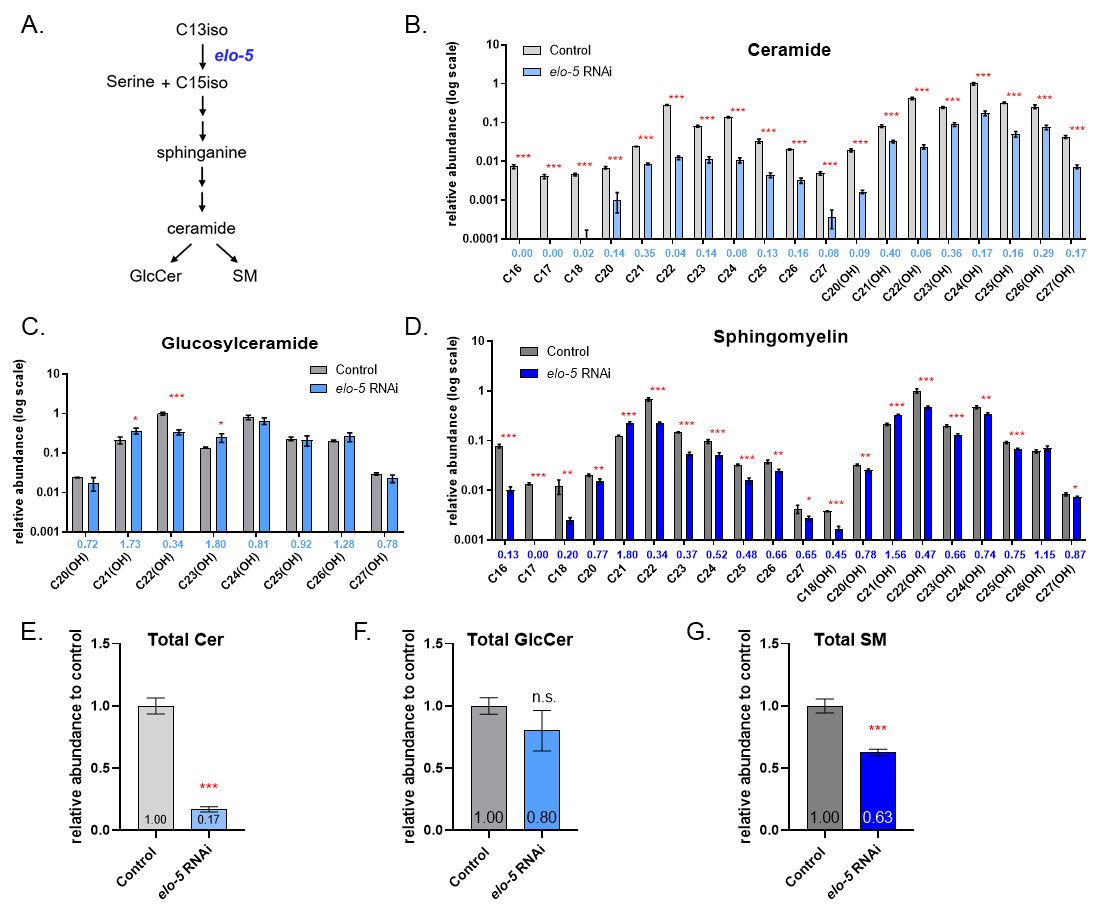

**Supplemental Figure S3. RNAi of each of the three ceramide synthase genes in *C. elegans* generated distinct GlcCer and SM profiles.**

(A) A heatmap showing abundance changes of individual glucosylceramide species in *hyl-1, hyl-2, lagr-1* RNAi worms relative to the control RNAi group. The average value of the three control RNAi samples, against which others were normalized, is set to 1. Abundance increase (fold change > 1) and decrease (fold change < 1) are indicated, respectively, by red and blue hues of varying saturation. (B-D) Relative abundance changes of individual GlcCer species and total GlcCer level changes in *hyl-1* (B)*, hyl-2* (C)*, and lagr-1* (D) RNAi worms. *n* = 3 biological replicates, data are shown as mean ± standard deviation (∗, *P* < 0.05; ∗∗, *P* < 0.01; ∗∗∗, *P* < 0.001). The colored values below each bar graph indicate the fold change of abundance of the RNAi-treated group versus the control group. (E) A heatmap showing abundance changes of individual sphingomyelin species in *hyl-1, hyl-2, lagr-1* RNAi worms relative to the control RNAi group. (F-H) Relative abundance changes of individual SM species in *hyl-1* (F)*, hyl-2* (G)*, and lagr-1* (H) RNAi worms. (I) Total SM level changes in *hyl-1, hyl-2, lagr-1* RNAi worms relative to the control RNAi worms.

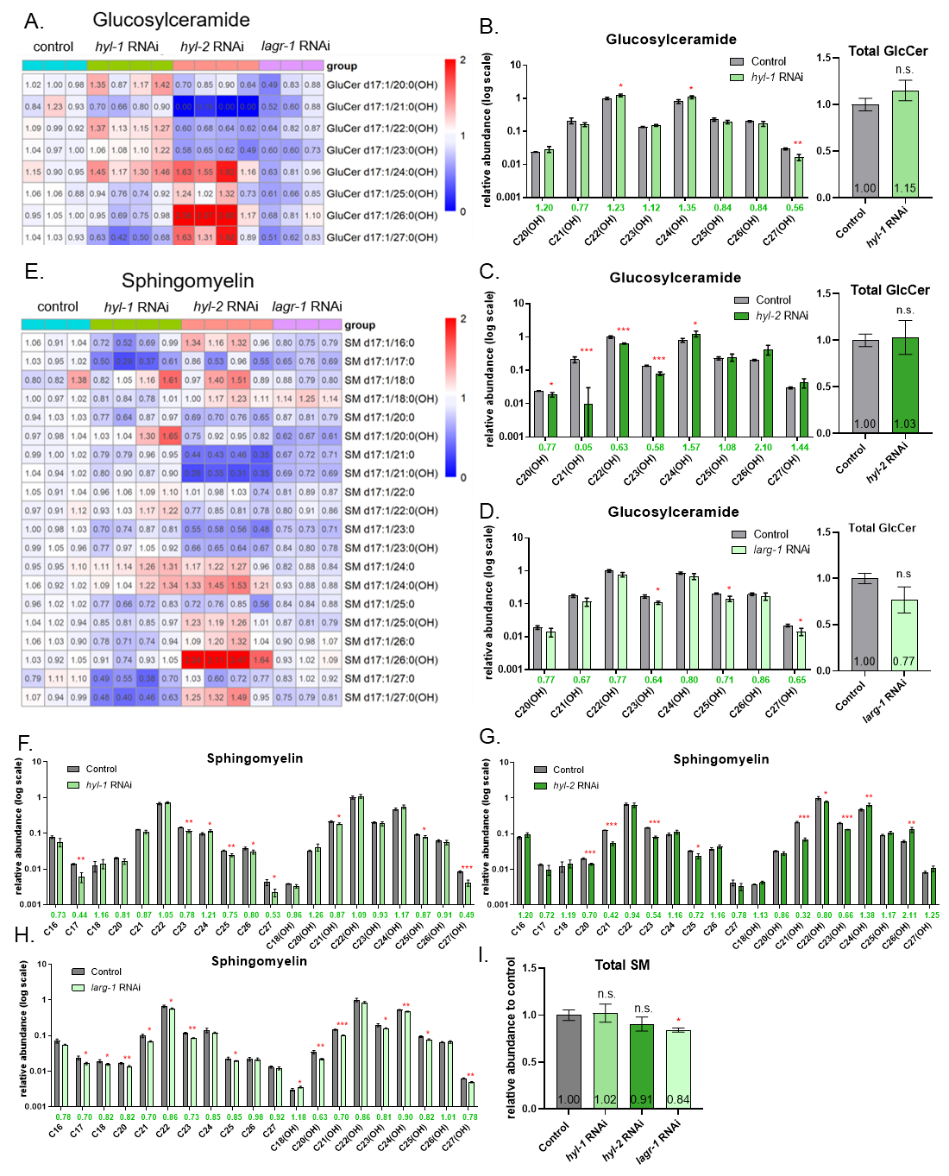

**Supplemental Figure 4. Cer and SM profiles of *C. elegans* treated with *cgt-1* or *cgt-3* RNAi**

(A-B) Relative abundance changes of individual Cer species in *cgt-1* (A) *and cgt-3* (B) RNAi worms. *n* = 3 biological replicates, data are shown as mean ± standard deviation (∗, *P* < 0.05; ∗∗, *P* < 0.01; ∗∗∗, *P* < 0.001). The colored values below each bar graph indicate the fold change of abundance of the RNAi-treated group versus the control group. (C) Total Cer level changes in *cgt-1 and cgt-3* RNAi worms relative to the control RNAi worms. (D-E) Relative abundance changes of individual SM species in *cgt-1* (D) *and cgt-3* (E) RNAi worms. (F) Total SM level changes in *cgt-1 and cgt-3* RNAi worms relative to the control RNAi worms.

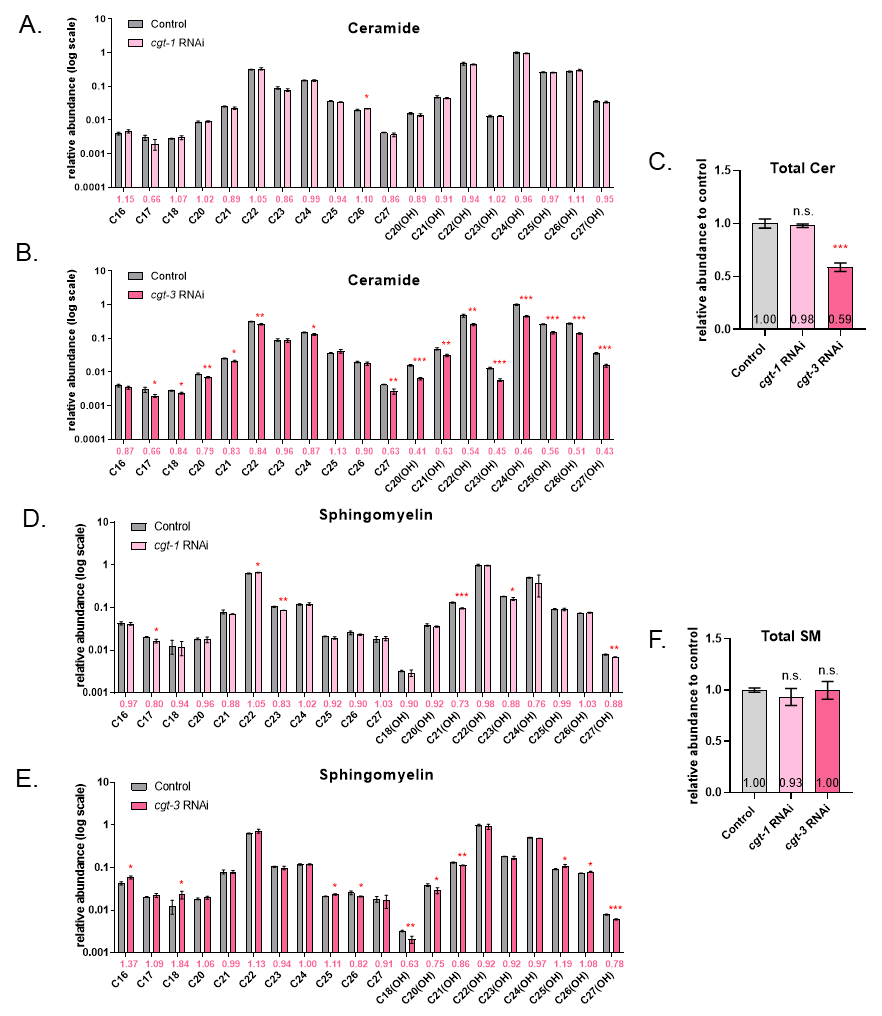

**Supplemental Figure S5. Cer and GlcCer profiles of *C. elegans* after RNAi of *sms-1, -2,* or *-3***

(A) A heatmap showing abundance changes of individual ceramide species in *sms-1, sms-2, sms-3* RNAi worms relative to the control RNAi group. The average value of the three control RNAi samples, against which others were normalized, is set to 1. Abundance increase (fold change > 1) and decrease (fold change < 1) are indicated, respectively, by red and blue hues of varying saturation. (B-D) Relative abundance changes of individual Cer species in *sms-1* (B)*, sms-2* (C)*, and sms-3* (D) RNAi worms. *n* = 2 or 3 biological replicates, data are shown as mean ± standard deviation (∗, *P* < 0.05; ∗∗, *P* < 0.01; ∗∗∗, *P* < 0.001). The colored values below each bar graph indicate the fold change of abundance of the RNAi-treated group versus the control group. (E) Total SM levels in *sms-1, sms-2, sms-3* RNAi worms relative to the control RNAi worms. (F) A heatmap showing abundance changes of individual GlcCer species in *sms-1, sms-2, sms-3* RNAi worms relative to the control RNAi group. (G-I) Relative abundance changes of individual GlcCer species and total GlcCer level changes in *sms-1* (G)*, sms-2* (H)*, and sms-3* (I) RNAi worms.

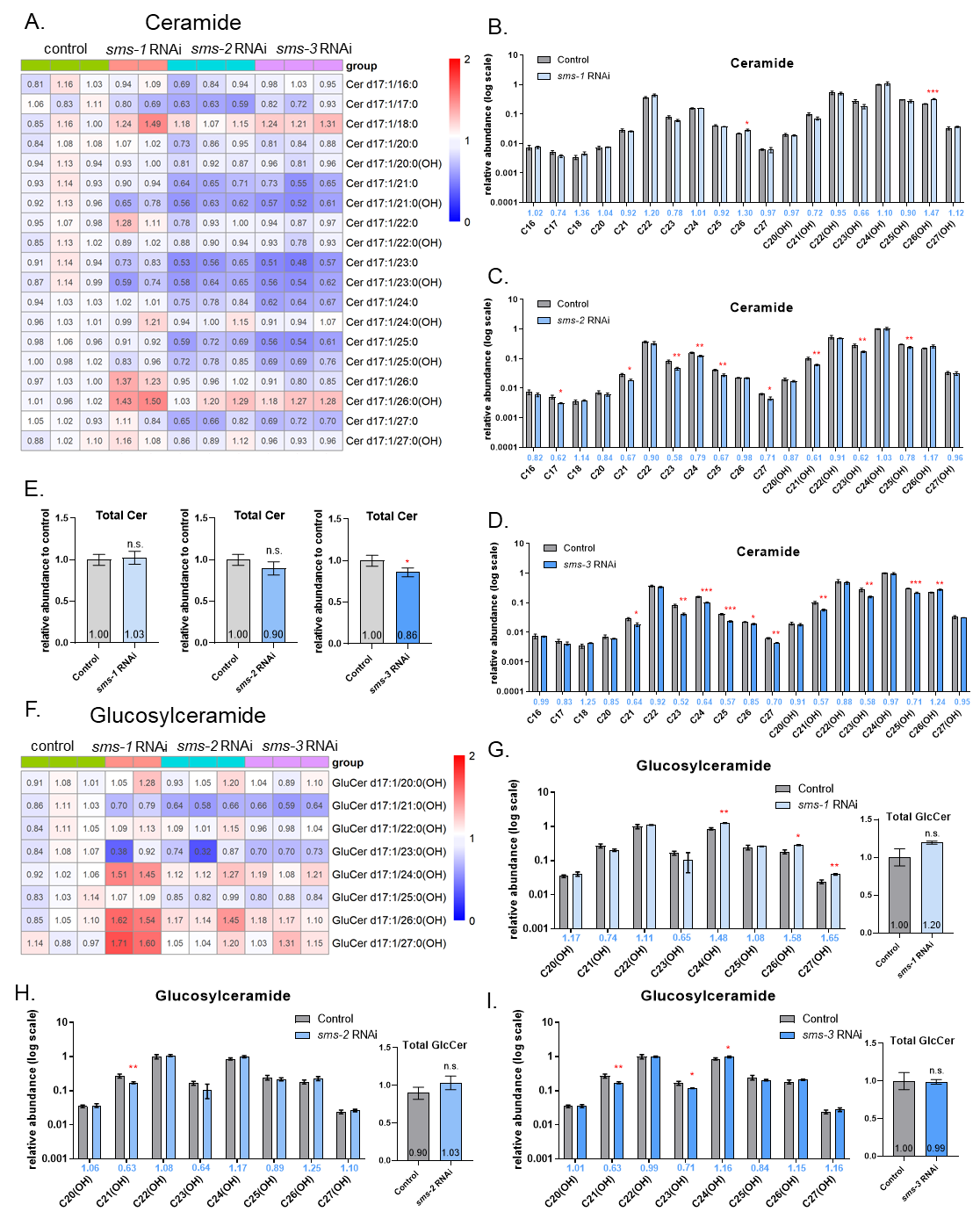

**Supplemental Figure S6. Relatedness in the relative abundance changes of different sphingolipids across ten RNAi conditions**

(A) A clustering plot of sphingolipid changes across all examined RNAi conditions. Red and blue indicate abundance increase and decrease, respectively. (B) The Pearson’s correlation coefficient values between sphingolipid species across all examined RNAi conditions. Red and blue denote positive and negative correlation, respectively.

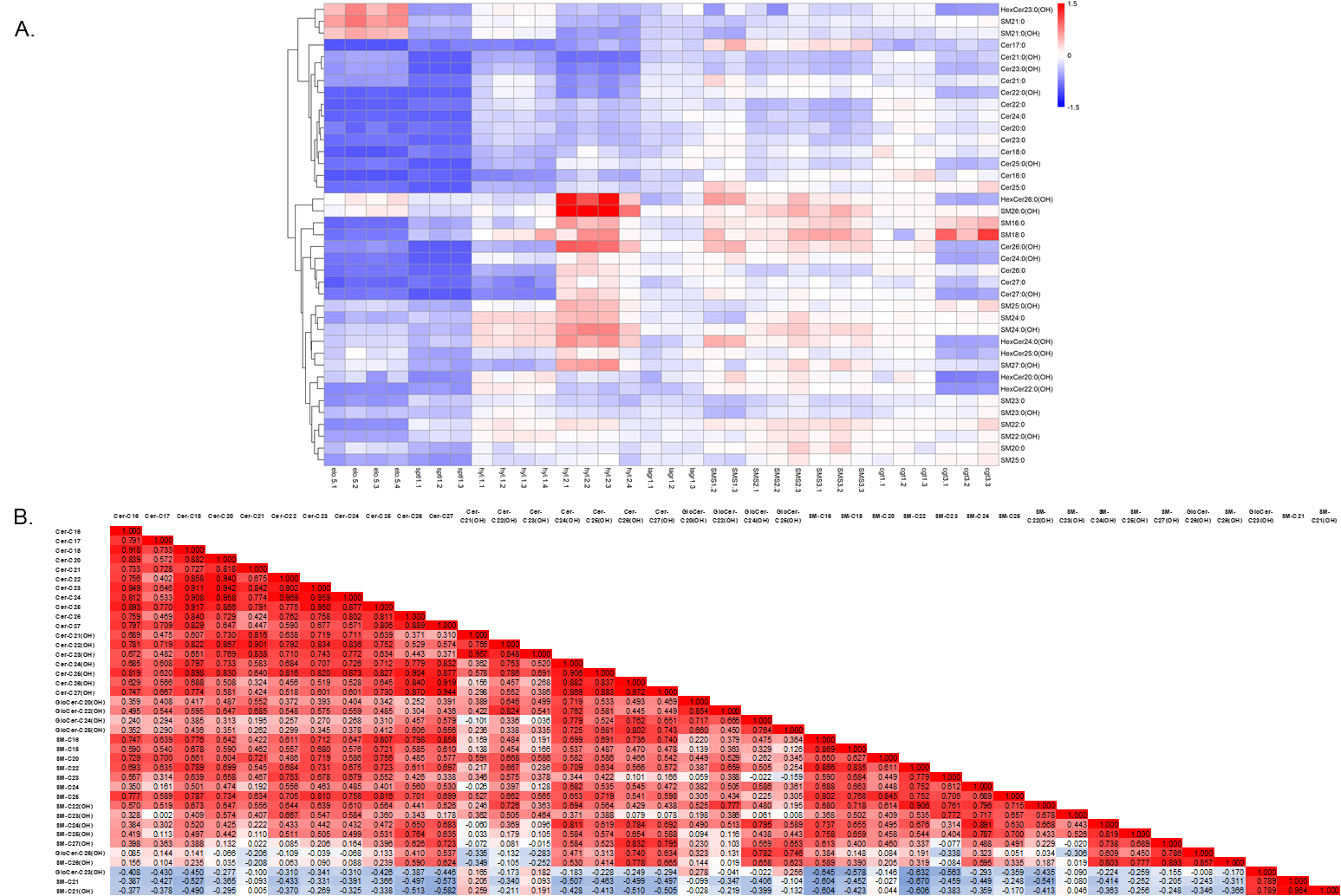
